# Uhrf1 governs the proliferation and differentiation of muscle satellite cells

**DOI:** 10.1101/2021.04.08.439096

**Authors:** Hiroshi Sakai, Yuichiro Sawada, Naohito Tokunaga, So Nakagawa, Iori Sakakibara, Yusuke Ono, So-ichiro Fukada, Tadahiko Kikugawa, Takashi Saika, Yuuki Imai

## Abstract

DNA methylation is an essential form of epigenetic regulation responsible for cellular identity. In muscle stem cells, termed satellite cells, DNA methylation patterns are tightly regulated during differentiation. However, it is unclear how these DNA methylation patterns are maintained. We demonstrate that a key epigenetic regulator, ubiquitin like with PHD and RING finger domains 1 (Uhrf1), is activated in proliferating myogenic cells but not expressed in quiescent or differentiated myogenic cells in mice. Ablation of Uhrf1 in mouse satellite cells impairs their proliferation and differentiation, leading to failed muscle regeneration. Loss of Uhrf1 in satellite cells alters transcriptional programs, leading to DNA hypomethylation with activation of Cdkn1a and Notch signaling. Although down-regulation of Cdkn1a rescued proliferation but not differentiation, inhibition of Notch signaling rescued impaired differentiation of Uhrf1-deficient satellite cells. These findings point to Uhrf1 as a regulator of self-renewal and differentiation of satellite cells via genome-wide DNA methylation patterning.

## Introduction

Adult skeletal muscle has a great capacity for regeneration, which is mediated by muscle satellite (stem) cells that express the *Pax7* gene (Seale et al., 2000). Genetic ablation of Pax7-expressing satellite cells in mice led to a drastic loss of muscle tissue and loss of muscle regeneration (Lepper et al., 2011; McCarthy et al., 2011; Murphy et al., 2011; Sambasivan et al., 2011). During regeneration, satellite cells are rapidly activated to proliferate and fuse to form new skeletal muscle fibers. Although it has been reported that many intrinsic and extrinsic cell signals contribute to the maintenance, activation, and differentiation of satellite cells, little information is available on the role of epigenetic regulation, especially maintenance of DNA methylation patterns, in satellite cells during muscle regeneration.

DNA methylation is critical for regulating gene transcription. Several studies reported that DNA methylation patterns are altered during the proliferation and differentiation of mouse and human myogenic cells (Carrió et al., 2015; Miyata et al., 2015; Tsumagari et al., 2013). A family of DNA methyltransferases (DNMT) mediate DNA methylation by enzymatic activities including *de novo* methylation (catalyzed by DNMT3a and DNMT3b) or maintenance of methylation (by DNMT1). Deleting DNMT3a in satellite cells led to loss of proliferation with increased expression of the *Cdkn1c* gene (Naito et al., 2016). Further, we reported that deletion of *Dnmt1* in satellite cells impaired muscle regeneration *in vivo*, resulting in a reduced number of satellite cells *in vitro* (Iio et al., 2021). However, it remains largely unknown how the DNA methylation pattern of satellite cells is maintained during proliferation and differentiation.

Ubiquitin like with PHD and RING finger domains 1 (Uhrf1; also known as Np95 in mice and ICB90 in humans) acts as an epigenetic regulator; it is essential for maintenance of DNA methylation by recruiting DNMT1 to hemi-methylated DNA sites (Bostick et al., 2007; Sharif et al., 2007), and it also can interact with DNMT3a and DNMT3b (Meilinger et al., 2009). Consistently, UHRF1 is up-regulated in proliferating cells, such as neural stem and basal stem cells in the airway, and down-regulated during differentiation or quiescence (Ramesh et al., 2016; Xiang et al., 2017). Previous reports have shown that ablation of *Uhrf1* in different cell types, including embryonic stem cells (Sharif et al., 2007), T cells (Obata et al., 2014), hematopoietic stem cells (Zhao et al., 2017), chondrocytes (Yamashita et al., 2017), and adult neural stem cells (Blanchart et al., 2018), leads to DNA hypomethylation, resulting in abrogation of proliferation and/or differentiation. However, no information is available on the functions of *Uhrf1* in satellite cells in muscle regeneration.

In this study, we demonstrate activation of *Uhrf1* in mouse satellite cells during regeneration. Deletion of *Uhrf1* in satellite cells led to impaired muscle regeneration with loss of satellite cells in mice. *Uhrf1*-deficient satellite cells exhibited dramatic changes in transcript levels and genome-wide DNA methylation profiles, which led to up-regulation of Cdkn1a and genes involved in Notch signaling. Therefore, we show that *Uhrf1* is critical for controlling the proliferation and differentiation of satellite cells via DNA methylation.

## Results

### Uhrf1 is active in satellite cells after muscle injury and during *in vitro* culture

To determine whether and when *Uhrf1* is expressed in muscles, the tibialis anterior (TA) muscles were injured by cardiotoxin (CTX) injection; uninjured and injured TA muscles were collected on different days post injury (dpi). During muscle regeneration, the *Uhrf1* transcript level peaked at 4 dpi and then decreased by 14 dpi (Figure S1A). To determine which myogenic cells contain active Uhrf1, immunofluorescence staining of Uhrf1 and Pax7 was performed in TA muscle sections (Figure 1A). In uninjured TA muscles, almost none of the Pax7+ satellite cells expressed Uhrf1 (Figure 1B). At 5 dpi, when the number of Pax7+ cells peaked during normal regeneration post-injury (Murphy et al., 2011), 11% of Pax7+ cells expressed Uhrf1, and this percentage decreased to 5.8% at 14 dpi (Figure 1B), which was compatible with the Uhrf1 transcript level in injured TA muscles. In addition, *Uhrf1* expression gradually decreased after induction of differentiation in C2C12 cells (Figures S1B–D).

**Figure 1.**
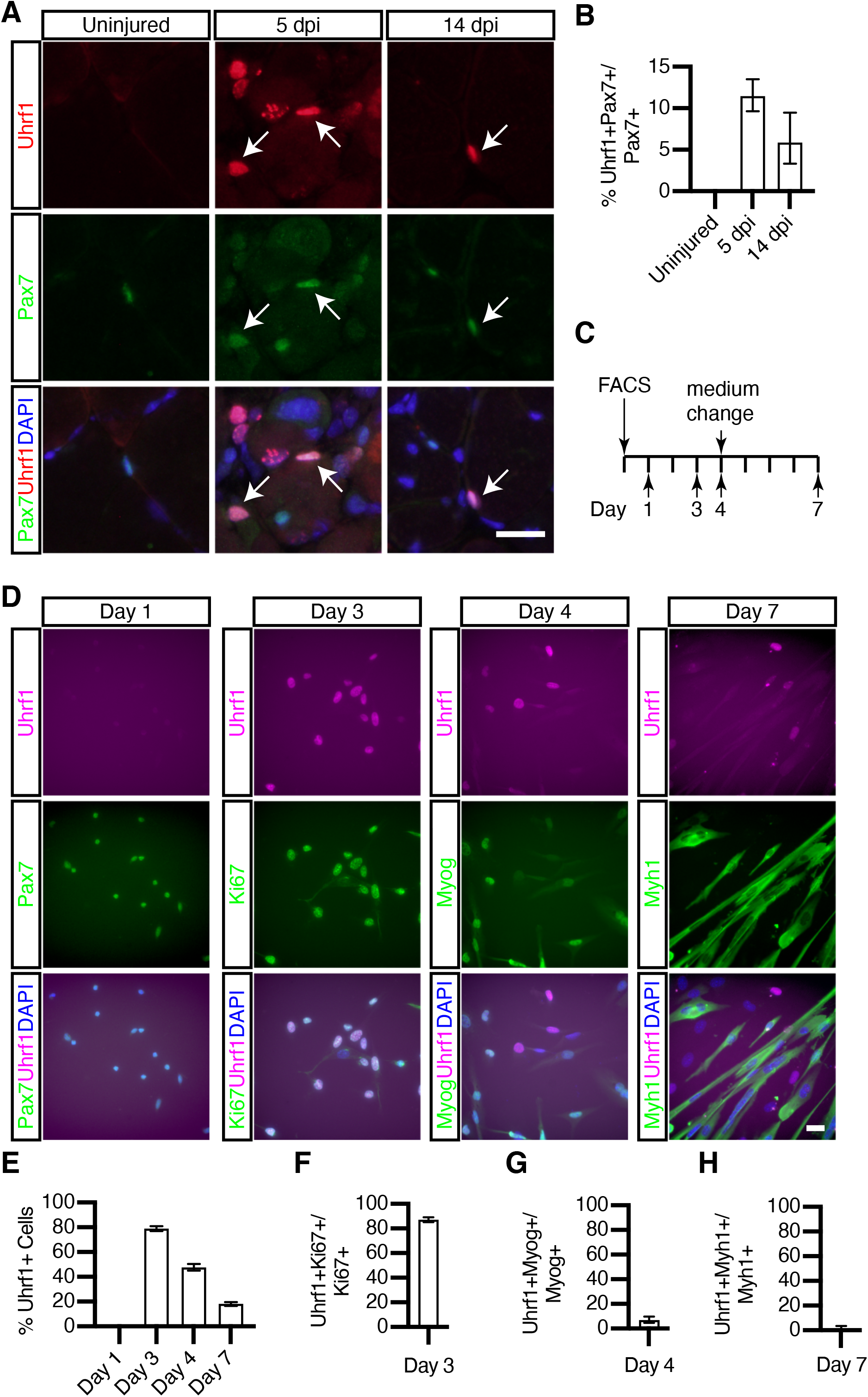
Uhrf1 is active in satellite cells after muscle injury and during *in vitro* culture. (A) Immunostaining of Pax7 and Uhrf1 during muscle regeneration. Arrows show Uhrf1+Pax7+ cells. (B) Quantification of Uhrf1+ cells among Pax7+ cells in uninjured and injured TA muscles collected at 5 and 14 dpi. (C) Time-course of satellite cells cultured *in vitro*. (D) Immunostaining of Uhrf1, Pax7, Ki67, Myogenin, and Myh1 after 1, 3, 4, and 7 days of culture. (E) Quantification of Uhrf1+ cells *in vitro*. (F) Quantification of Uhrf1+ cells among proliferating (Ki67+) cells after 3 days of culture. (G) Quantification of Uhrf1+ cells among Myogenin+ cells after 4 days of culture. (H) Quantification of Uhrf1+ cells in Myh1+ fibers after 7 days of culture. For all graphs, data were pooled from three independent experiments and are expressed as the mean with 95% CI. Scale bars = 20 µm. See also Figure S1.

To further examine the expression pattern of Uhrf1, satellite cells were isolated by FACS and cultured for up to 7 days *in vitro* (Figure 1C). Almost all cells were positive for Pax7 on day 1, but none of these cells expressed Uhrf1 (Figure 1D), which supported the *in vivo* results in intact muscles. The majority of myogenic cells (78%) were positive for Uhrf1 on day 3, and this percentage gradually decreased during differentiation (Figure 1E). Whereas 87% of the Uhrf1+ cells expressed Ki67 on day 3 (Figure 1F), only 6.9% of Myogenin+ cells (differentiated) expressed Uhrf1 on day 4 (Figure 1G), and almost no Myh1+ myotubes expressed Uhrf1 (Figure 1H). These observations indicate that Uhrf1 is not expressed in satellite cells during quiescence but is transiently expressed in proliferating myogenic cells before being down-regulated during differentiation.

### Loss of Uhrf1 in satellite cells impairs muscle regeneration

To determine whether *Uhrf1* expression in satellite cells is necessary for muscle regeneration, we selectively deleted this gene using *Pax7*^*CE/+*^ (Lepper et al., 2009) and *Uhrf1*^*fl/fl*^ (Skarnes et al., 2011) mice. The efficiency of *Uhrf1* ablation in satellite cells in *Pax7*^*CE/+*^*;Uhrf1*^*fl/fl*^ mice was measured by isolating satellite cells by FACS from hindlimb muscles of *Pax7*^*CE/+*^*;Uhrf1*^*fl/fl*^ mice following administration of oil (control mice) or tamoxifen (TMX, mutant mice) for 5 consecutive days (Figures S2A and S2B). Deletion was highly efficient as the *Uhrf1* transcript level was 90% lower in mutant mice than control mice (Figure S2C). Uhrf1 protein was expressed in 65% of satellite cells cultured for 3 days from control mice but in 5.2% of those from mutant mice (Figures S2D and S2E). Together, these experiments indicate that Uhrf1 expression was effectively down-regulated in satellite cells from *Pax7*^*CE/+*^*;Uhrf1*^*fl/fl*^ mice.

To investigate the effect of Uhrf1 expression loss in satellite cells on muscle regeneration, the TA muscles of control and mutant mice were injured by CTX and harvested at 5 dpi (Figure 2A). As *Pax7*^*CE/+*^ mice treated with TMX (Cre control mice) exhibit a mild muscle regeneration phenotype (Mademtzoglou et al., 2018; Sakai et al., 2020), Cre control mice were included in the experiments for regeneration. The regenerating fibers, characterized by expression of embryonic myosin heavy chain (Myh) 3, were detected at 5 dpi in control mice (Figure 2B), similar to a previous report (Murphy et al., 2011). Although there was no difference in the mass (Figure S2F) or cross-sectional area (CSA) of TA muscles (Figure S2G), the Myh3 level was decreased in Cre control mice compared with the control, while the proportion of Myh3+ regenerating fibers was severely reduced in mutant mice (Figure 2C). These results indicate that loss of *Uhrf1* affects the regeneration of new fibers. In addition, there was a significant decrease in the number (Figure 2E) and proliferation of Pax7+ cells (Figure 2F) at 5 dpi in mutant mice compared with control and Cre control mice.

**Figure 2.**
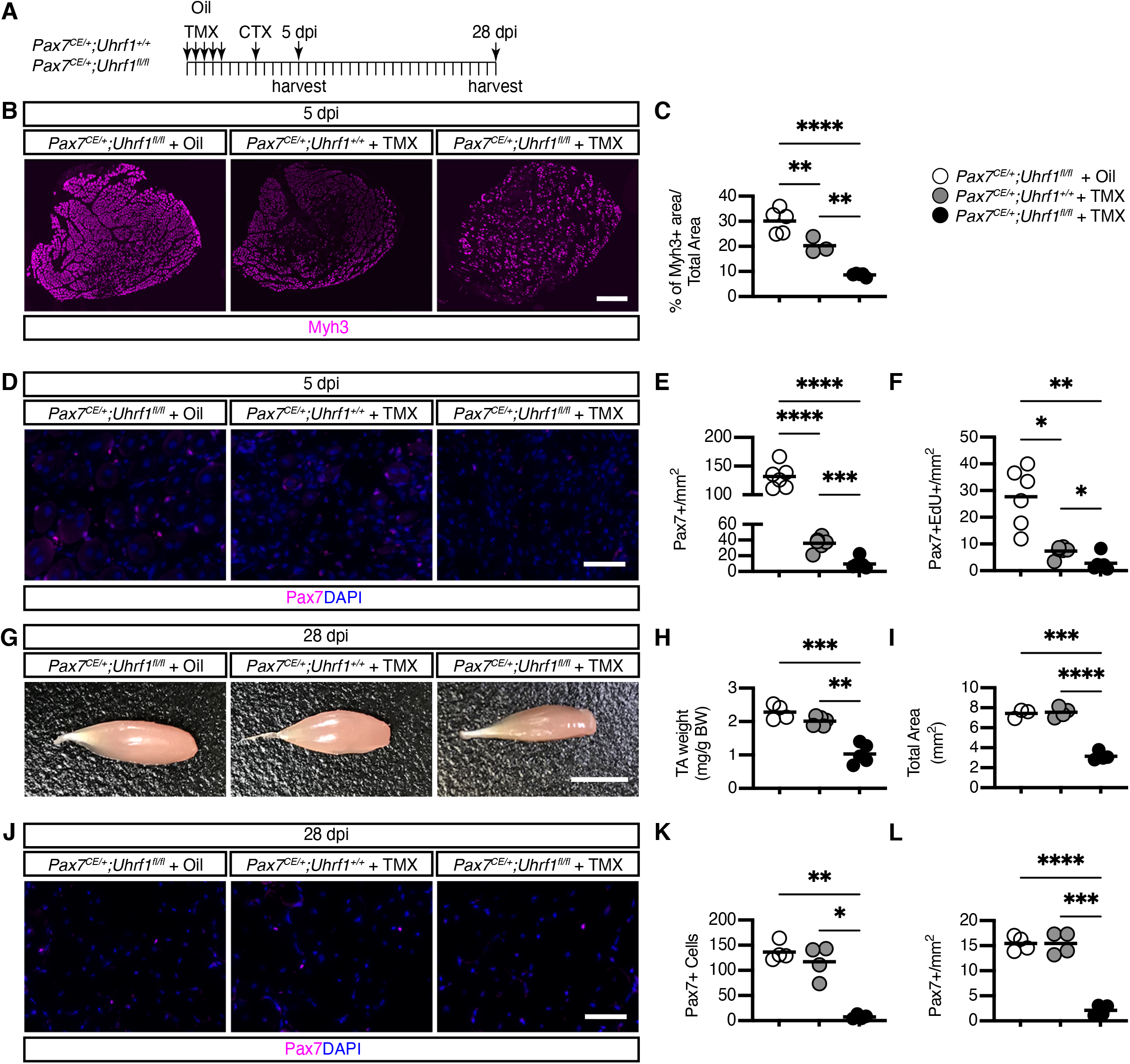
Loss of Uhrf1 expression in satellite cells impairs muscle regeneration. (A) Experimental design for TMX treatment followed by CTX injection and sample harvest at 5 and 28 dpi. (B) Immunostaining of Myh3 in TA muscles at 5 dpi. (C) The average area of Myh3+ regenerating myofibers in TA muscle cross-sections at 5 dpi (n = 3 or 5 mice/condition). (D) Immunostaining of Pax7 in TA muscle cross-sections at 5 dpi. (E) Quantification of Pax7+ cells at 5 dpi (n = 6 mice/condition). (F) Quantification of Pax7+EdU+ cells at 5 dpi (n = 6 mice/condition). (G) Whole mount TA at 28 dpi. (H) Mass of TA muscles at 28 dpi (n = 4 or 5 mice/condition). (I) The average total area of TA muscle cross-sections at 28 dpi (n = 3 or 4 mice/condition). (J) Immunostaining of Pax7 in TA muscle cross-sections at 28 dpi. (K, L) The total and average number of Pax7+ cells in TA muscle cross-sections at 28 dpi (n = 4 mice/condition). \**p* < 0.05, \*\**p* < 0.01, \*\*\**p* < 0.001, \*\*\*\**p* < 0.0001. Scale bars: 50 µm in (D and J), 500 µm in (B), and 50 mm in (G). See also Figure S2.

To determine whether the observed phenotype at 5 dpi was sustained during the muscle regenerative process, injured TA muscles were harvested at 28 dpi (Figure 2A), at which time regeneration is largely complete (Murphy et al., 2011). Although there was no difference in the TA muscle mass between control and Cre control muscle according to macroscopic observations (Figure 2G), the mutant mice exhibited significant muscle mass loss in the TA muscles (Figure 2H). There was a significant decrease in the total CSA in mutant mice (Figures 2I and S2H) but no difference between the control and Cre control mice. Also, TA sections contained a significantly lower number of Pax7+ cells in mutant mice compared with control mice (Figure 2K), and this significant decrease remained even after normalization to the total CSA (Figure 2L). In summary, loss of Uhrf1 expression in satellite cells significantly impaired proliferation and muscle regeneration.

To investigate the mechanisms of satellite cell impairment in *Pax7*^*CE/+*^*;Uhrf1*^*fl/fl*^ mice, single myofibers were isolated from the extensor digitorum longus muscle of control and mutant mice and cultured for up to 4 days (Figure S2I). On days 0 and 2, the number of Pax7+ satellite cells per myofiber was comparable between the mutant and control mice (Figure S2J). In contrast, the total number of satellite cells and the proportion of Pax7-MyoD+ cells, which are committed to activation and differentiation, were decreased in the myofibers of mutant mice compared with control mice (Figure S2J).

To further characterize the impaired proliferation and differentiation observed in culture, satellite cells from control and mutant mice were characterized *in vitro* (Figure S2K). In EdU labelling experiments to assess proliferation, approximately 70% of satellite cells were positive for EdU in control mice, compared with 33% in mutant mice, on day 3 (Figures S2L and S2M). Furthermore, the number of Myogenin+ cells was significantly decreased in mutant mice on day 4 (Figures S2N and S2O). In addition, myotube formation, observed by Myh1 staining on day 7, was also compromised in the mutant mouse cells (Figures S2P and S2Q). Taken together, these *in vitro* findings indicate that Uhrf1 is required in myogenic cells for proliferation but not activation, and the loss of Uhrf1 leads to impairments of differentiation.

### Loss of Uhrf1 in satellite cells alters DNA methylation patterns and subsequently transcriptional programs

To identify the genes regulated by Uhrf1 in satellite cells, RNA-Seq was performed in satellite cells obtained from control and mutant mice (Figure 3A). A heatmap of the expression profiles showed a clear distinction between the control and mutant satellite cells (Figure S3A). We detected 2044 up-regulated and 1833 down-regulated genes in the mutant compared with control satellite cells (Figures 3B and S3B). According to Gene Ontology (GO) enrichment analysis, the down-regulated genes showed significant enrichment of the biological process terms cell proliferation and cell cycle (Figure 3C), consistent with the impairment of these processes observed in the mutant cells. The genes that were up-regulated in mutant satellite cells showed enrichment of GO terms associated with “vasculature development”. In addition, several gene sets associated with impaired cell proliferation or differentiation, such as “positive regulation of proteolysis” and “apoptotic signaling pathway”, were also enriched among the up-regulated genes (Figure 3C). However, only caspase 9 in the critical apoptotic signaling pathway was up-regulated in the mutant cells (Table 1). These results suggest that activation of apoptosis may not be a main cause of the phenotype of the mutant satellite cells.

**Table 1.** Analysis of senescence- and apoptosis-related gene expression by RaNA-Seq.

| **Symbol** | **ExpMea**<br>**n** | **Exp**<br>**Control** | **Exp Mutant** | **log2FC** | **p-value** | **padj** |
| --- | --- | --- | --- | --- | --- | --- |
| Cdkn1a | 600.15 | 473.807 | 726.494 | 0.616 | 4.8e-5 | 6.0e-4 |
| Cdkn1b | 25.838 | 27.355 | 24.321 | −0.163 | 0.064 | 0.242 |
| Cdkn2b | 9.607 | 8.206 | 11.007 | 0.383 | 0.079 | 0.283 |
| Cdkn2a | 18.511 | 17.514 | 19.509 | 0.148 | 0.487 | 0.85 |
| Trp53 | 165.366 | 163.689 | 167.043 | 0.029 | 0.8 | 0.926 |
| Casp9 | 5.16 | 4.78 | 5.54 | 0.178 | 0.004 | 0.029 |
| Casp6 | 13.381 | 12.815 | 13.946 | 0.114 | 0.024 | 0.114 |
| Casp8 | 18.767 | 17.74 | 19.794 | 0.15 | 0.217 | 0.555 |
| Casp7 | 7.432 | 7.88 | 6.984 | −0.154 | 0.288 | 0.656 |
| Casp3 | 36.873 | 36.342 | 37.404 | 0.04 | 0.599 | 0.917 |

**Figure 3.**
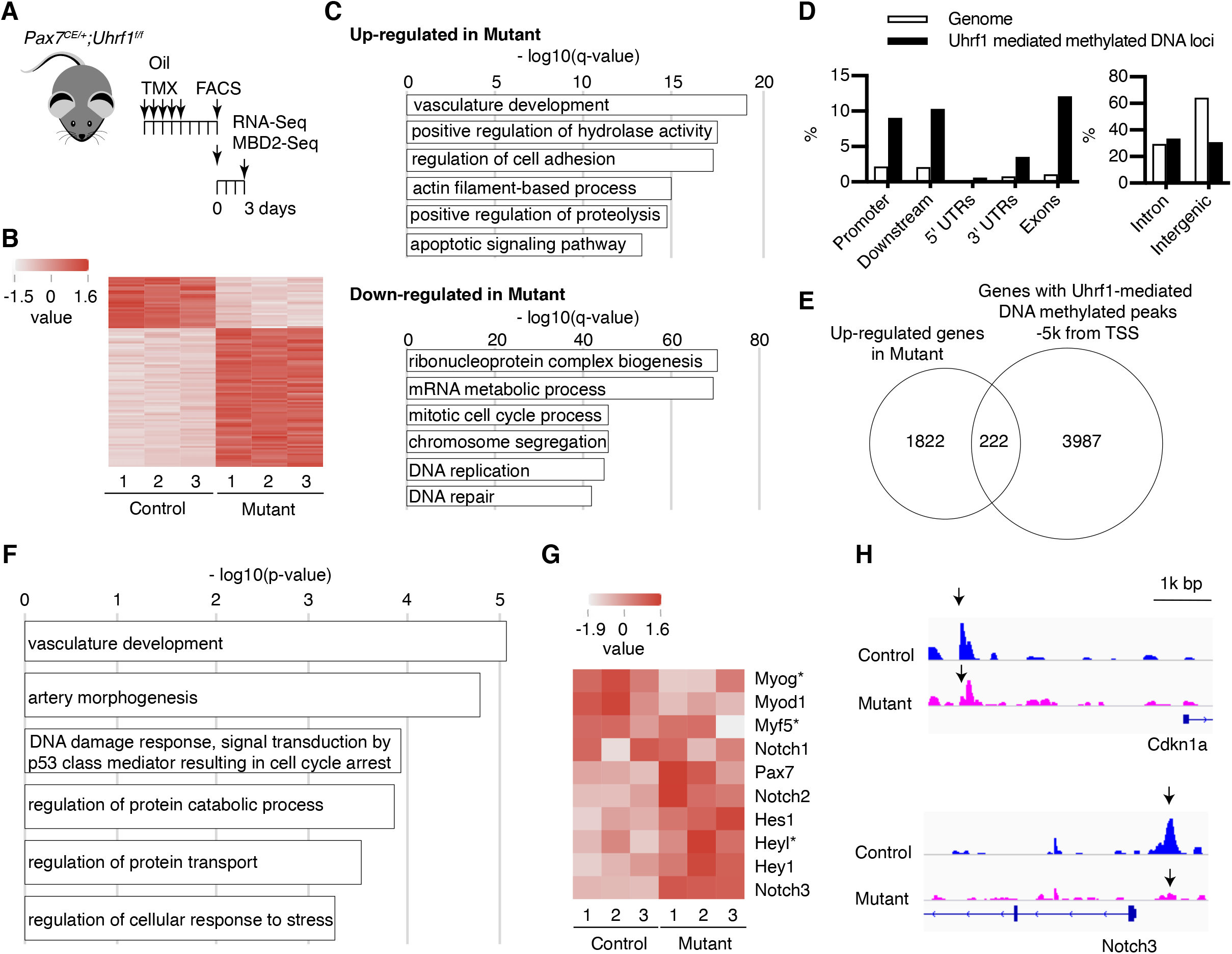
Loss of Uhrf1 in satellite cells alters DNA methylation patterns and thereby transcriptional programs. (A) Experimental design for RNA-Seq and MBD2-Seq. (B) Heatmap of the expression values (normalized as transcripts per million, TPMs) of the top 100 differentially expressed genes in each sample. (C) Bar graph of the Gene Ontology (GO) terms enriched among the up-regulated and down-regulated genes in mutant satellite cells. (D) Distribution of Uhrf1-mediated methylated DNA with the given intervals and scores, with the genomic features determined using a cis-regulatory element annotation system. (E) Venn diagram of the up-regulated genes associated with Uhrf1-mediated methylated DNA loci. (F) Bar graph of the GO terms enriched among the up-regulated genes associated with Uhrf1-mediated methylated DNA loci. (G) RNA-Seq analyses of Notch and myogenic-related genes (normalized as TPMs) in each sample. Genes with an asterisk were not differentially expressed. (H) Methylated DNA signals around the *Cdkn1a* and *Notch3* loci in control and mutant satellite cells visualized using IGV. See also Figure S3.

To determine whether the genes up-regulated by Uhrf1 deficiency were specific to satellite cells, we compared the up-regulated genes in chondrocytes (Yamashita et al., 2017) and hematopoietic stem cells lacking *Uhrf1* (Zhao et al., 2017) with those in satellite cells (Figure S3C). Among the 2044 genes up-regulated in Uhrf1 deficient-satellite cells, 331 overlapped with the 1122 genes up-regulated in *Uhrf1*-knockout chondrocytes. On the other hand, only 77 genes overlapped with the 626 genes up-regulated in hematopoietic stem cells lacking *Uhrf1*. These results indicate that *Uhrf1* deficiency induces up-regulation of partially overlapped genes, but cell-specific knockout of Uhrf1 may lead to up-regulation of cell-specific genes.

Most DNA methylation occurs at transposable elements (TEs) within the mouse genome. Loss of DNA methylation due to Uhrf1 deficiency in embryonic stem cells (Sharif et al., 2007) or neural stem cells (Ramesh et al., 2016) leads to activation of TEs. We thus examined the expression of TEs using RNA-Seq data using STAR program (v2.7.6a) (Dobin et al., 2013) with dedicated parameters and algorithms for TE analysis (see Materials and Methods). As expected, several TEs were up-regulated in the mutant satellite cells, but almost none were down-regulated (Figure S3D). We found up-regulation of intracisternal A particles (IAPs) in mutant compared with control satellite cells (Figure S3E), which is in line with the results from *Uhrf1*-deficient embryonic stem cells (Bostick et al., 2007; Sharif et al., 2007).

We next performed methyl-CpG binding domain protein 2-enriched genome sequencing (MBD2-Seq) in the control and mutant satellite cells (Figure 3A). The higher GC count per read in methylated DNA compared with unmethylated DNA (Figure S3F) and the distinct methylation patterns observed at the *H19* and *Kcnq1ot1* loci, which are imprinted genes (Figure S3G), indicated proper enrichment of methylated DNA by MBD2. To identify DNA loci methylated by Uhrf1, peak calling was performed using MACS2. We identified 39,231 peaks, and these loci were enriched in promoters, downstream regions, and gene bodies of the genome (Figure 3D). This change in the DNA methylation profile might not be a secondary result of decreased Dnmt expression, as the expression of *Dnmt1, Dnmt3a*, and *Dnmt3b* did not change drastically compared with *Uhrf1* deficiency in mutant satellite cells (Figure S3H), as determined by RNA-Seq. To identify Uhrf1 target genes in satellite cells, we analyzed the genes up-regulated by *Uhrf1* deficiency that overlapped with the Uhrf1-mediated DNA methylated peaks. We found 222 genes with Uhrf1-mediated peaks within -5000bp from the transcriptional start site (Figure 3E). GO enrichment analysis showed that these genes are associated with “vasculature development” and “cell cycle arrest”, which is consistent with the results of the GO enrichment analysis of the RNA-Seq data (Figure 3F). Among these genes, *Cdkn1a* was a candidate mediator of “cell cycle arrest” (Table 1), consistent with the reduced proliferation of the mutant satellite cells. We also found that genes annotated as “vasculature development” included those related to Notch signaling that play critical roles in sustaining the quiescent state of satellite cells (Fukada et al., 2011), and their expression was up-regulated in the mutant satellite cells (Figure 3G). We also detected reduced DNA methylation levels in the regions upstream of the *Cdkn1a* and *Notch3* loci in mutant satellite cells (Figure 3H). These results suggest that *Uhrf1* deficiency alters DNA methylation patterns, followed by transcriptional changes in satellite cells.

### Suppression of Cdkn1a and Notch signaling rescued the impaired proliferation and differentiation of Uhrf1-deficient satellite cells

To explore whether up-regulation of the Cdkn1a or Notch signaling pathway is responsible for the impaired proliferation and differentiation of *Uhrf1*-deficient satellite cells, we treated mutant satellite cells with siRNA targeting *Cdkn1a* or with DAPT, an inhibitor of γ-secretase (Figure 4A). Although Cre control satellite cells showed up-regulation of *Cdkn1a* (Figure S4A) and reduced proliferation compared with control mice, siRNA knockdown of *Cdkn1a* in mutant satellite cells (Figure S4A) significantly rescued the impaired proliferation of the mutant satellite cells (Figures 4B and 4C). However, *Cdkn1a* knockdown was not sufficient to rescue the impaired differentiation of the mutant satellite cells (Figures 4D and 4E). In contrast, although DAPT treatment did not affect the proportion of EdU+ cells (Figures 4F and 4G), unlike siRNA-mediated knockdown of *Cdkn1a*, it partially rescued proliferation in terms of absolute cell numbers (Figure S4C), and DAPT treatment suppressed HeyL expression (Figure S4B) and rescued differentiation, as measured by Myogenin expression (Figures 4H and 4I), in mutant satellite cells. We also confirmed that knockdown of *Cdkn1a* (Figures S4D and S4E) and DAPT treatment (Figures S4F and S4G) did not selectively expand the population of surviving Uhrf1+ cells among mutant satellite cells. Taken together, Uhrf1-mediated DNA methylation negatively regulated the expression of Cdkn1a and Notch signaling for physiological proliferation and/or differentiation of satellite cells.

**Figure 4.**
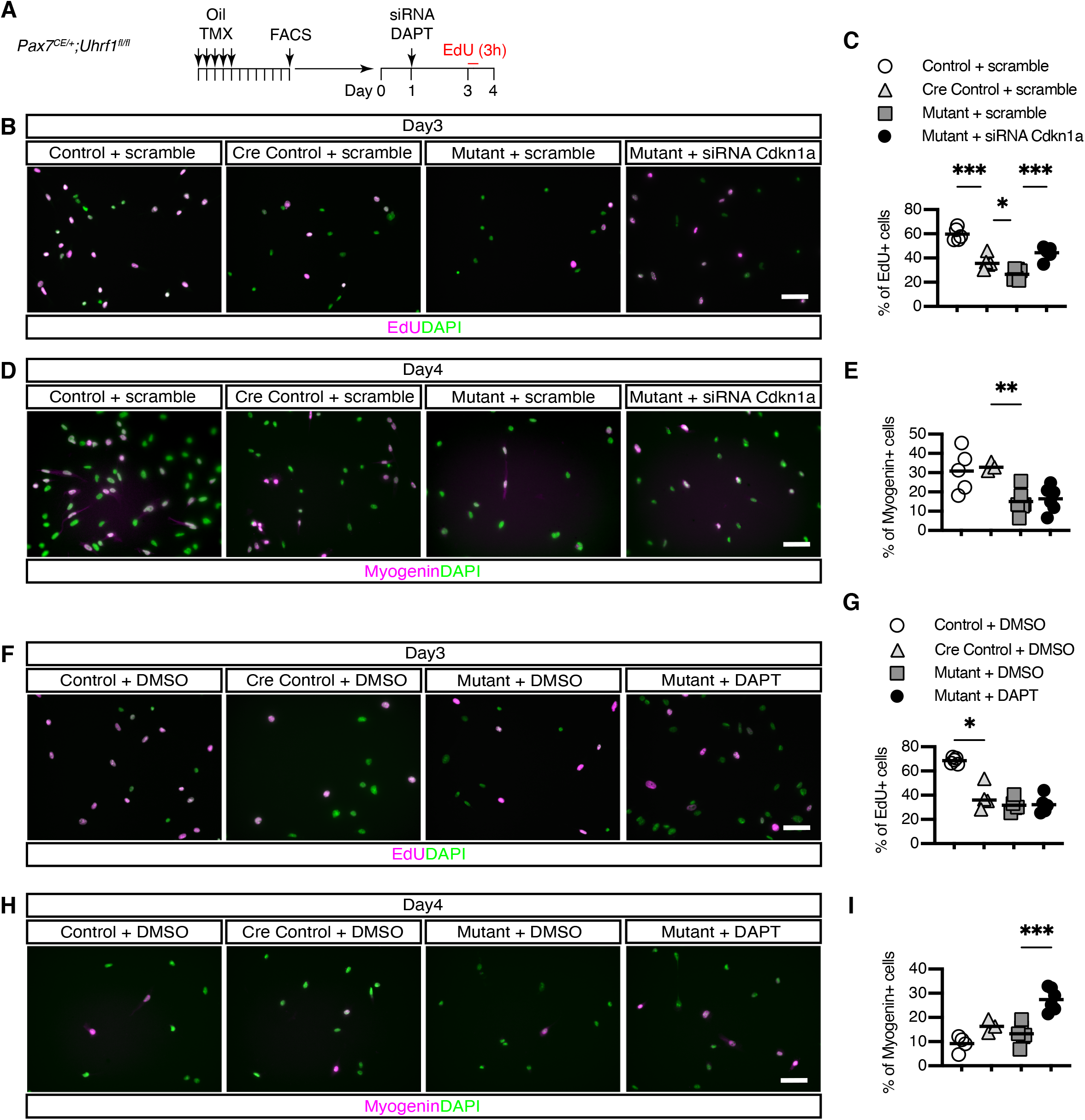
Decreased Cdkn1a and Notch signaling rescues the impaired proliferation and differentiation of Uhrf1-deficienct satellite cells. (A) Experimental design for the siRNA and DAPT treatments. (B) EdU detection in *Cdkn1a*-knockdown satellite cells. (C) Quantification of EdU+ *Cdkn1a*-knockdown satellite cells (n = 5 for control and Cre control; n = 6 for mutant cells). (D) Immunostaining of Myogenin in *Cdkn1a*-knockdown satellite cells. (E) Quantification of Myogenin+ *Cdkn1a*-knockdown satellite cells (n = 5 for control; n = 3 for Cre control; n = 6 for mutant cells). (F) EdU detection in satellite cells treated with DAPT. (G) Quantification of EdU+ DAPT-treated satellite cells (n = 5 for control and mutant; n = 4 for Cre control cells). (H) Immunostaining of Myogenin in satellite cells treated with DAPT. (I) Quantification of Myogenin+ DAPT-treated satellite cells (n = 4 for control; n = 3 for Cre control; n = 6 for mutant cells). \**p* < 0.05, \*\**p* < 0.01, \*\*\**p* < 0.001. Scale bars = 50 µm. See also Figure S4.

## Discussion

In this study, we identified *Uhrf1* as a critical regulator of skeletal muscle satellite cells. Notably, *Uhrf1* is not expressed during quiescence but is up-regulated during proliferation and down-regulated during differentiation in satellite cells. Our findings agree with previous reports showing that Uhrf1 is necessary for maintaining the DNA methylation profile during DNA replication in proliferating cells. Studies on neural stem cells and basal stem cells in airways showed that *Uhrf1* is up-regulated during proliferation and down-regulated during differentiation or quiescence (Ramesh et al., 2016; Xiang et al., 2017). As *Uhrf1* overexpression was reported to contribute to cancer initiation and progression (Mudbhary et al., 2014), tight regulation of *Uhrf1* expression in stem cells appears to be critical for self-renewal and differentiation.

Further, ablation of Uhrf1 in satellite cells led to impaired muscle regeneration, similar to Dnmt1- (Iio et al., 2021) or Dnmt3a-deficient mice (Naito et al., 2016). The marked loss of EdU incorporation indicated a remarkable reduction in the number of *Uhrf1*-deficient satellite cells *in vivo* and *in vitro* at 28 dpi. Therefore, the impairment of muscle regeneration of mutant mice could result from decreased numbers of myogenic cells.

In addition, we report that loss of *Uhrf1* in satellite cells alters transcriptional programs caused by alteration of genome-wide DNA methylation patterns. Consistent with the impaired proliferation of *Uhrf1*-deficient satellite cells, we found that many cell-cycle-related genes were down-regulated. We propose that Uhrf1 deficiency in mutant satellite cells results directly in hypomethylation and general up-regulation of transcription. Indeed, the GO term most enriched among the up-regulated genes by hypomethylation in *Uhrf1*-deficient satellite cells was “vascular development”. As satellite cells have a potential to adopt a pericyte phenotype following stimulation of the Notch and PDGF signaling pathways (Gerli et al., 2019), loss of Uhrf1 in satellite cells might trigger partial transdifferentiation into vascular-like cells. We found that TEs, especially IAPs, were strongly up-regulated in *Uhrf1*-deficient satellite cells. This phenotype is comparable with that of *Uhrf1*-deficient cortical neural stem cells showing strong activation of IAPs (Ramesh et al., 2016). Since long-terminal repeats in IAPs could change gene regulation at a given locus (Morgan et al., 1999), further studies are needed to clarify whether transposition of IAPs into new genomic loci would contribute to the phenotype of mutant satellite cells.

Finally, we showed that suppression of the increased Cdkn1a and Notch signaling in *Uhrf1*-deficienct satellite cells rescued their impaired proliferation and differentiation. Cdkn1a was identified as a target of Uhrf1 in neural stem cells and colonic regulatory T cells (Blanchart et al., 2018; Obata et al., 2014). Thus, loss of *Uhrf1* might lead to cell cycle arrest in diverse cell types. We found that silencing of *Cdkn1a* in mutant satellite cells rescued their decreased cell proliferation, but not differentiation, suggesting that *Uhrf1*-mediated satellite cells require additional factors for differentiation. Notch signaling plays a critical role in sustaining the quiescent state of satellite cells. Notch3 is highly expressed in quiescent satellite cells (Fukada et al., 2007), and its expression is strongly up-regulated by overexpression of the Notch intracellular domain (Wen et al., 2012). These results suggest that cell autonomous Notch signaling inhibits muscle differentiation of mutant satellite cells. Thus, we propose that Uhrf1 epigenetically regulates, at least in part, satellite cell proliferation via Cdkn1a-mediated cell cycle regulation, and loss of Uhrf1 in myogenic cells leads to up-regulation of Notch signaling and thereby decreased differentiation and proliferation of satellite cells.

In conclusion, the present study demonstrates that Uhrf1 is a critical epigenetic regulator of proliferation and differentiation in myogenic cells, by controlling gene expression via maintenance of DNA methylation.

## Acknowledgments

We thank S. Tajbakhsh for critically reading the manuscript, S. Nakanishi and A. Nishio for their technical support, K. Kameda for the flow cytometry support and other members of the Division of Analytical Bio-Medicine and the Division of Laboratory Animal Research, the Advanced Research Support Center, Ehime University. This work was supported in part by MEXT/JSPS KAKENHI (JP18H06439 and JP19K19947 to H.S.; JP23689066, JP15H04961, JP15K15552, JP17K19728, JP19H03786 to Y.I.); HIRAKU-Global Program, which is funded by MEXT’s “Strategic Professional Development Program for Young Researchers” (to H.S.); and The Nakatomi Foundation and Takeda Science Foundation (to H.S. and Y.I.).

## Author Contributions

Conceptualization, H.S. and Y.I. Methodology, H.S., S.N., I.S., Y.O., and S.F. Software, H.S., N.T., and S.N. Validation, H.S. and Y.S. Formal analysis, H.S., N.T., and S.N. Investigation, H.S., Y.S., and N.T. Resources, Y.O. and S.F. Data curation, H.S. and N.T. Writing – original draft, H.S. and Y.I. Writing – review & editing, H.S. and Y.I. Visualization, H.S. and Y.S. Supervision, H.S., T.K., T.S., and Y.I. Project administration, Y.I. Funding acquisition, H.S., T.K., T.S., and Y.I.

## Methods

### Animals

C57BL/6JJcl mice were purchased from CLEA Japan. The *Pax7*^*CE/+*^ (Lepper et al., 2009; Jackson Laboratories, stock no: 012476) and *Uhrf1*^*fl/fl*^ (*B6Dnk;B6N-Uhrf1*^*<tm1a(EUCOMM)Wtsi>/*^*Ieg*; Skarnes et al., 2011; European Mouse Mutant Archive) mouse strains have been described previously. *Uhrf1*^*fl/fl*^ mice were crossed with *Pax7*^*CE/+*^ mice to generate *Pax7*^*CE/+*^*;Uhrf1*^*fl/fl*^ mice. All mice were housed in a specific pathogen-free facility under climate-controlled conditions and a 12-h light/dark cycle and were provided water and a standard diet ad libitum. All male animals were evaluated at age 7–12 weeks old. Tamoxifen (150 µl, 20 mg/ml) (Sigma, cat# T5648) dissolved in corn oil (Sigma, cat# C8267) was injected intraperitoneally for 5 consecutive days to induce Cre-mediated recombination.

### Muscle regeneration

Injury was induced by injecting 100 µl 10 µM cardiotoxin (CTX) solution (Latoxan, cat# L8102) into the left tibialis anterior (TA) muscles under anesthesia using isoflurane. The right TA served as the intact control.

### Immunofluorescence and microscopy

For immunofluorescence (IF) staining, muscles were frozen in liquid nitrogen-chilled isopentane, and 10 µm cryosections were prepared. For Pax7 and Uhrf1 staining, sections were fixed in 4% paraformaldehyde (PFA) in PBS for 5 min and then boiled in antigen retrieval buffer (10 mM sodium citrate, pH 6) for 10 min. For Myh3 staining, tissue sections were fixed for 10 min in acetone at −20°C before boiling, blocked in 5% goat serum (Gibco, cat# 16210-064) in PBS for 60 min, and incubated with the primary antibodies overnight at 4°C. The antibodies used for IF staining were anti-Pax7 (DSHB, cat# PAX7, RRID:AB_528428; 1/5 dilution; or Invitrogen, cat# PA1-117, RRID:AB_2539886; 1/100 dilution), anti-Uhrf1 (Santa Cruz Biotechnology, cat# sc-373750, RRID: AB_10947236; 1/50 dilution), anti-Myh3 (DSHB, cat# F1.652, RRID:AB_528358; 1/20 dilution), Alexa fluor 568 goat anti-mouse IgG1 (Invitrogen, cat# A-21124; 1/1000 dilution), and Alexa fluor 488 goat anti-rabbit IgG (Invitrogen, cat# A-11008; 1/1000 dilution).

For IF staining in cells, cells were fixed in 4% PFA in PBS for 10 min, followed by permeabilization using 0.5% Triton X-100 in PBS for 10 min. Cells were blocked using 5% goat serum in PBS. The following primary antibodies were used: anti-Uhrf1 (Santa Cruz Biotechnology, cat# sc-373750, RRID: AB_10947236; 1/100 dilution; or MBL, cat# D289-3 RRID:AB_10597260; 1/100 dilution), anti-Ki67 (Novus, cat# NB500-170, RRID:AB_10001977; 1/100 dilution), anti-Myogenin (Santa Cruz Biotechnology, cat# sc-12732, RRID:AB_627980; 1/100 dilution), and anti-Myh1 (DSHB, cat# MF20, RRID:AB_2147781; 1/10 dilution). The signals were visualized using the following fluorochrome-coupled secondary antibodies: Alexa fluor 568 goat anti-mouse IgG1a (Invitrogen, cat# A-21124), Alexa fluor 488 goat anti-rabbit IgG (Invitrogen, cat# A-11008), and Alexa fluor 488 goat anti-rat IgG (Invitrogen, cat# A-11006). Nuclei were counterstained with DAPI.

Stained cells or tissues were photographed using BIOREVO (Keyence) and quantified using Fiji (https://imagej.net/Fiji). The numbers of Uhrf1+, Pax7+, Ki67+, Myogenin+, and EdU+ cells *in vitro* and the area of Myh3+ fibers *in vivo* were determined using the Fiji Analyse Particles function. For total muscle CSA quantification, the whole area of transverse TA muscle sections was measured by laminin staining. The numbers of Uhrf1+, Pax7+, and EdU+ cells *in vivo* were determined by manual counting using the Adobe Photoshop count tool.

### Isolation of satellite cells

Isolation of satellite cells from hindlimb muscles was performed as described previously (Liu et al., 2015) with minor modifications. Briefly, muscles were chopped in cold PBS and digested using 800 U/ml collagenase type 2 (Worthington, cat# LS004177) in F10 medium (Gibco, cat# 11550-043) supplemented with horse serum (Gibco, cat# 26050-088) and antibiotic–antimycotic (Gibco, cat# 15240-062) for 60 min followed by another digestion using 1000 U/ml collagenase type 2 and 11 U/ml dispase (Gibco, cat# 17105-041) for 30 min. The supernatants were filtered through a 40 µm cell strainer (Falcon, cat# 352340) and centrifuged to yield a cell suspension. The following antibodies were used: CD31-FITC (BioLegend, cat# 102406; RRID:AB_312901; 1/100 dilution), CD45-FITC (BioLegend, cat# 103108, RRID:AB_312973; 1/100 dilution), Sca1-APC (BioLegend, cat# 122512, RRID:AB_756197; 1/100 dilution), and Vcam1-PE (Invitrogen, cat# 12-1061-82, RRID:AB_2572573; 1/100 dilution). Cells were isolated using the FACS Aria II (Becton, Dickinson and Company) and analyzed by FlowJo (https://www.flowjo.com).

### Cell culture, siRNA transfection, and DAPT treatment

The C2C12 cell line was culture in DMEM (Gibco, cat# 10569-010) supplemented with 10% FBS (Gibco, cat# 10099-141) and antibiotic–antimycotic. The medium was replaced with differentiation medium (DMEM supplemented with 2% horse serum) to induce myotube formation.

Isolated satellite cells were resuspended in DMEM/F12 (Gibco, cat# 10565-018) supplemented with 20% FBS, 2% Ultroser G (Pall, cat# 15950-017), and antibiotic–antimycotic. Cells at 5000 or 10000/cm^2^ were cultured on an 8-well chamber slide (Thermo Scientific, cat# 177445) or in a 6-well plate coated with 1 mg/ml Matrigel (Corning, cat# 354230). Cells were transfected with non-targeting siRNA (Dharmacon, cat# D-001810-10-05) or siRNA targeting Cdkn1a (Dharmacon, cat# L-058636-00-0005) using Lipofectamine RNAiMAX (Thermo Scientific, cat# 13778-075) following the manufacturer’s protocol. DAPT (20 µM, Calbiochem, cat# 565784) in DMSO was used to inhibit Notch signaling.

### EdU labelling and detection

For *in vivo* labelling, EdU (5-ethynyl-2’-deoxyuridine, Invitrogen, cat# A10044) was dissolved in PBS at 0.5 µg/µl and injected intraperitoneally into mice at 5 µg/g body weight 24 h before harvesting. For *in vitro* labelling, the cells were incubated with EdU (10 µM) for 3 h. EdU was detected using Click-iT EdU Imaging Kits (Invitrogen, cat# C10340) with Alexa Flour 647.

### RNA isolation and quantitative real-time PCR

Total RNA was extracted from satellite cells using the RNeasy Plus Micro Kit (Qiagen, cat# 74034) following the manufacturer’s protocol. cDNA was synthesized from total RNA using PrimeScript (Takara, cat# RR036A). qPCR was performed in duplicate samples using TB Green Premix Ex Taq II (Takara, cat# RR820S) and Thermal Cycler Dice (Takara, cat# TP850). The primer sequences are listed in Table 2.

**Table 2.**
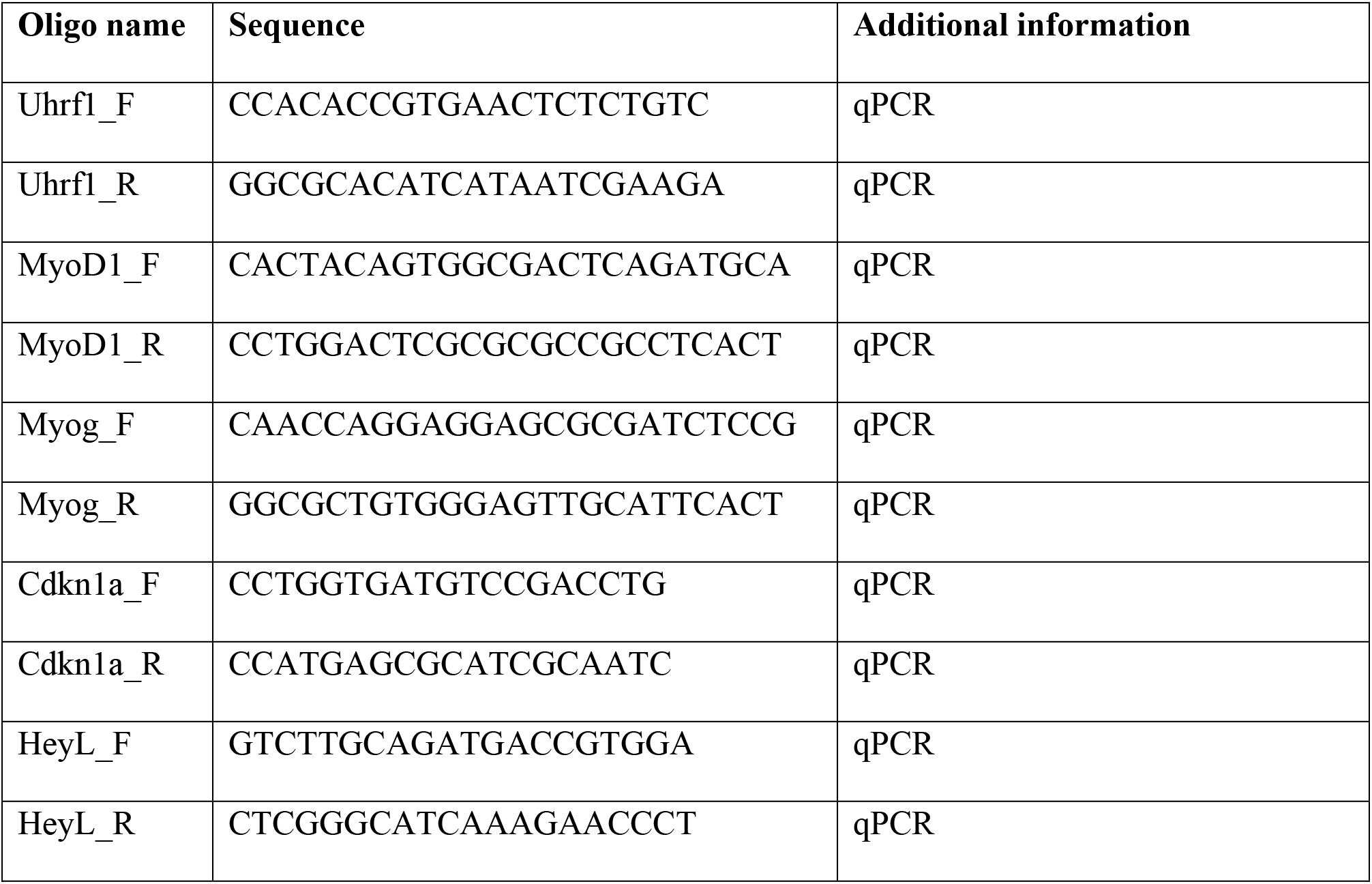
Oligonucleotides used in this study.

### Isolation, culture, and immunostaining of single myofibers

Single myofibers were isolated from extensor digitorum longus muscles as described previously (Rosenblatt et al., 1995). The isolated fibers were fixed in 4% PFA or cultured in DMEM supplemented with 30% FBS, 1% chicken embryo extract (US Biological, cat# C3999), and 10 ng/ml bFGF (Wako, cat# 062-06661). Immunostaining was performed following a previous protocol (Shinin et al., 2009).

### RNA-Seq and data analysis

The integrity of isolated RNA was verified using the Agilent 2100 Bioanalyzer with RNA 6000 Pico Kit (Agilent, cat# 5067-1513). RNA-Seq libraries were prepared using the NEBNext Ultra II Directional RNA Library Prep Kit for Illumina (NEB, cat# E7760S) with the NEBNext rRNA Depletion Kit (NEB, cat# E6310L) and NEBNext Multiplex Oligos for Illumina (NEB, cat# E7335S) according to the manufacturer’s instructions. The constructed libraries were verified using the Agilent 2100 Bioanalyzer with DNA 1000 kit (Agilent, cat# 5067-1504) and quantified using KAPA Library Quantification Kits (KAPA Bio, cat# KK4824) and the 7500 Real-Time PCR System (ABI, cat# 7500-01). The libraries were sequenced as 150 bp paired-end reads using the NovaSeq 6000 Sequencing System (Illumina). The expression analysis was performed using RaNA-Seq (Prieto and Barrios, 2019). Expression level was quantified by the Salmon algorithm (Patro et al., 2017) using the paired-end, default parameters. Differential expression analysis was performed by DESeq2 (Love et al., 2014) using the following parameters: test = Wald; fitType = local. GO enrichment analysis was performed using Metascape (Zhou et al., 2019).

For transposable element (TE) analysis, raw reads were trimmed using TrimGalore (v0.6.6; http://www.bioinformatics.babraham.ac.uk/projects/trim_galore/) and mapped to the reference genome (GRCm38, PRI) with annotation (PRI, https://www.gencodegenes.org/mouse/) using STAR (v2.7.6a) (Dobin et al., 2013). The option ‘--runMode alignReads --runThreadN 4 outMultimapperOrder Random --outSAMtype BAM SortedByCoordinate --outSAMmultNmax 1 -- outFilterMultimapNmax 1000 --outFilterMismatchNmax 3 --winAnchorMultimapNmax 1000 -- alignIntronMax 1 --alignMatesGapMax 350 --alignEndsType EndToEnd’ was used, allowing a higher number of multi-mapped reads (Teissandier et al., 2019). TE families were quantified using featureCounts (v2.0.1; -M -T 1 -s 0 -p) (Liao et al., 2014) and TE annotation (http://labshare.cshl.edu/shares/mhammelllab/www-data/TEtranscripts/TE_GTF/). The differentially expressed TEs were extracted by DESeq2.

### MBD2-Seq and analysis of the sequencing data

Genomic DNA was isolated and purified using the NucleoSpin Tissue Kit (MACHEREY-NAGEL, cat# 740952). Purified DNA was sheared using the Covaris S220 ultrasonicator with microTUBE AFA (Covaris, cat# 520045). Methylated and non-methylated DNA fragments were separated using the EpiXplore Methylated DNA Enrichment Kit (Clontech, cat# PT5034-2). Libraries were generated using the DNA SMART ChIP-Seq Kit (Takara, cat# 634865). The libraries were selected from 250–350-bp fragments obtained by E-Gel 2% SizeSelect electrophoresis (Invitrogen, cat# G661012). Size-selected libraries were verified using the Agilent 2100 Bioanalyzer with the High Sensitivity DNA kit (Agilent, cat# 5067-4626) and quantified using KAPA Library Quantification Kits (KAPA Bio, cat# KK4824) and the 7500 Real-Time PCR System (ABI, cat# 7500-01). The libraries were sequenced using NovaSeq 6000 with 150 bp paired end reads. Raw reads were down-sampled to 50 million reads for each condition to avoid the bias due to different sequencing depths. The reads were trimmed using TrimGalore and were mapped using HISAT2 (Kim et al., 2015).

Peaks were obtained by MACS2 (v2.2.6) (Zhang et al., 2008), using cells derived from TMX-treated mice as the ‘control,’ with the following options: --slocal 0 --llocal 0 -q 0.0001. ChIPpeakAnno (Zhu et al., 2010) was used to annotate and visualize the peaks. Methylated DNA signals were visualized using Integrative Genomics Viewer (IGV).

### Statistical analysis

Data were analyzed using Prism 9 (GraphPad Software). Two-sample test for equality of proportions was performed by prop.test in R (https://www.r-project.org). Welch’s t-test was used to compare variable between two groups. Welch’s ANOVA with Dunnett’s T3 multiple comparison test was used to compare variables among three groups.

### Data availability

The datasets generated from this study are available in Gene Expression Omnibus with accession number GSE169193.

### Study approval

All animals were maintained and used according to a protocol approved by the Animal Experiment Committee of Ehime University, Japan.

## Supplemental information titles and legends

**Supplementary Figure 1.**
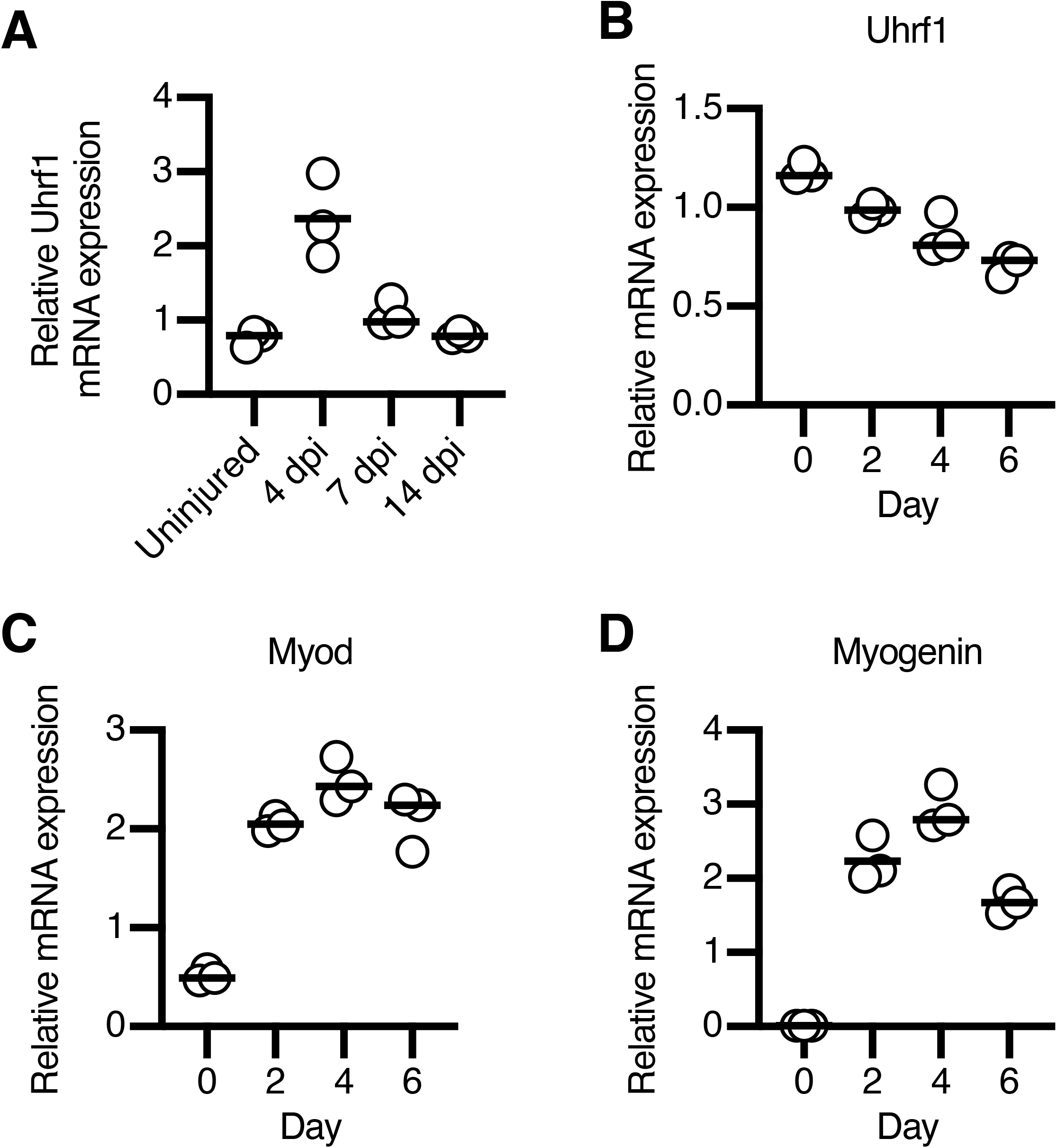
Uhrf1 is down-regulated during muscle differentiation in C2C12 cells. (A) RT-qPCR analysis of *Uhrf1* in uninjured and injured TA muscles (n = 3 mice for each time point). (B–D) RT-qPCR analysis of (B) *Uhrf1*, (C) *Myod*, and (D) *Myogenin* during C2C12 differentiation (n = 3 for each time point).

**Supplementary Figure 2.**
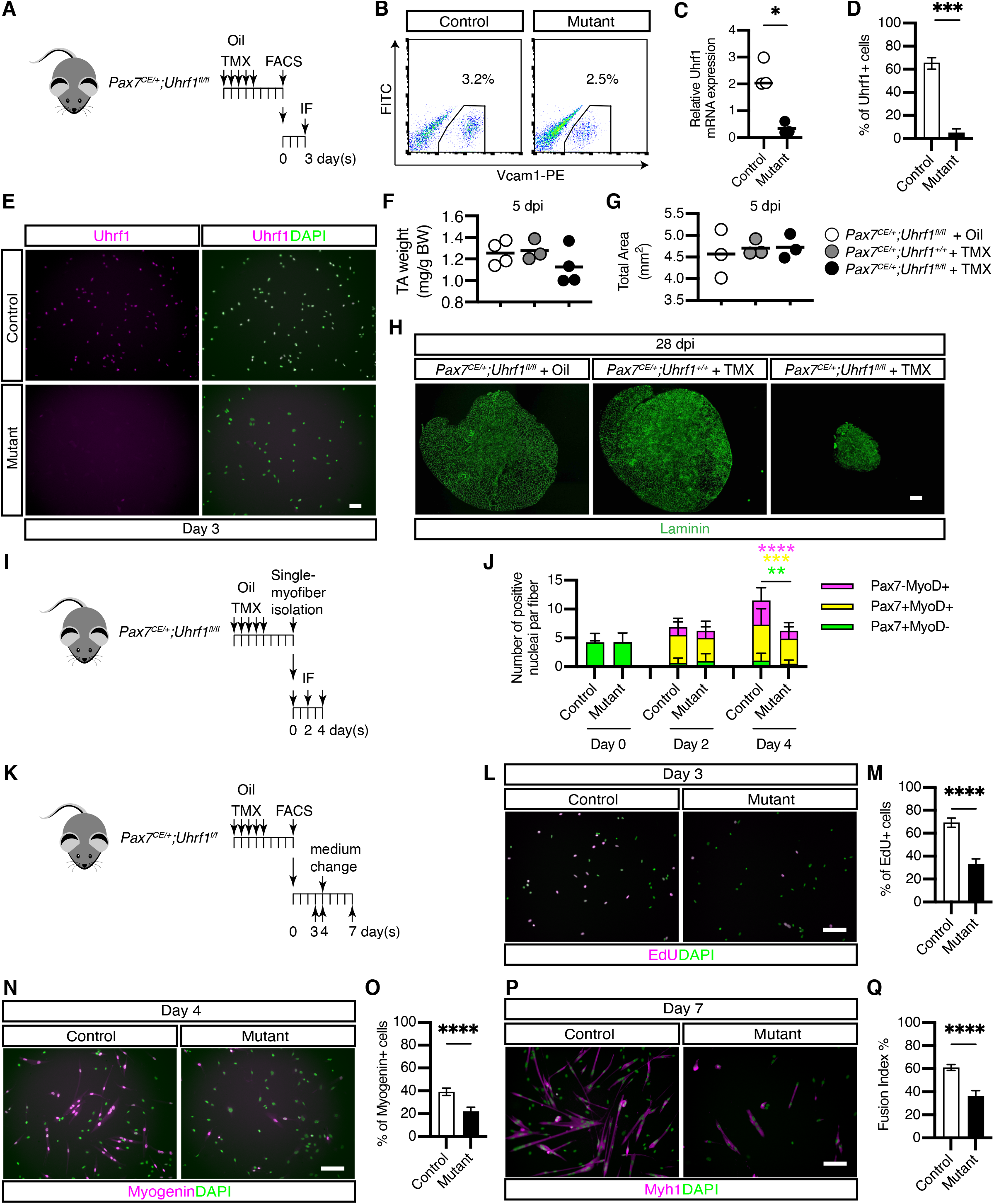
Deletion of Uhrf1 in satellite cells in *Pax7*^*CE/+*^*;Uhrf1*^*fl/fl*^ mice disturbs proliferation and differentiation of satellite cells. (A) Experimental design for TMX followed by FACS and immunostaining after 3 days in culture. (B) Representative FACS profiles of satellite cells from oil- or TMX-treated *Pax7*^*CE/+*^*;Uhrf1*^*fl/fl*^ mice. (C) RT-qPCR analysis of *Uhrf1* in satellite cells from oil- or TMX-treated *Pax7*^*CE/+*^*;Uhrf1*^*fl/fl*^ mice (n = 3 mice/condition). (D) Quantification of Uhrf1+ cells by IF (n = 362 cells for oil-treated mice, n = 304 cells for TMX-treated mice). Data are expressed as the mean with 95% CI. (E) Immunostaining of Uhrf1 in satellite cells from oil- or TMX-treated *Pax7*^*CE/+*^*;Uhrf1*^*fl/fl*^ mice. (F) Mass of TA muscles at 5 dpi (n = 3 or 4 mice/condition). (G) The average of total area in TA muscle cross-sections at 5 dpi (n = 3 mice/condition). (H) Immunostaining of laminin in TA at 28 dpi. (I) Experimental design for TMX followed by isolating single fibers. (J) Quantification of Pax7+ and Myod+ cells by immunostaining during myogenic differentiation in single fibers (n = 150-600 cells for each group, data represent the mean ± SD). (K) Experimental design for TMX followed by FACS and IF after 3, 4, and 7 days in culture. (L) EdU detection in satellite cells from oil- or TMX-treated *Pax7*^*CE/+*^*;Uhrf1*^*fl/fl*^ mice after 3 days in culture. (M) Quantification of EdU+ cells after 3 days in culture. (N) Immunostaining of Myogenin after 4 days in culture. (O) Quantification of Myogenin+ cells after 4 days in culture. (P) Immunostaining of Myh1 after 7 days in culture. (Q) Fusion index after 7 days in culture. For graphs in D, M, O, and Q, data were pooled from three independent experiments and data are expressed as the mean with 95% CI. Welch’s t-test (C and J), two-sample test for equality of proportions with continuity correction (D, M, O, and Q), \**p* < 0.05, \*\**p* < 0.01, \*\*\**p* < 0.001, \*\*\*\**p* < 0.0001. Scale bars = 50 µm in (E), 500 µm in (H), and 100 µm in (L), (N), and (P).

**Supplementary Figure 3.**
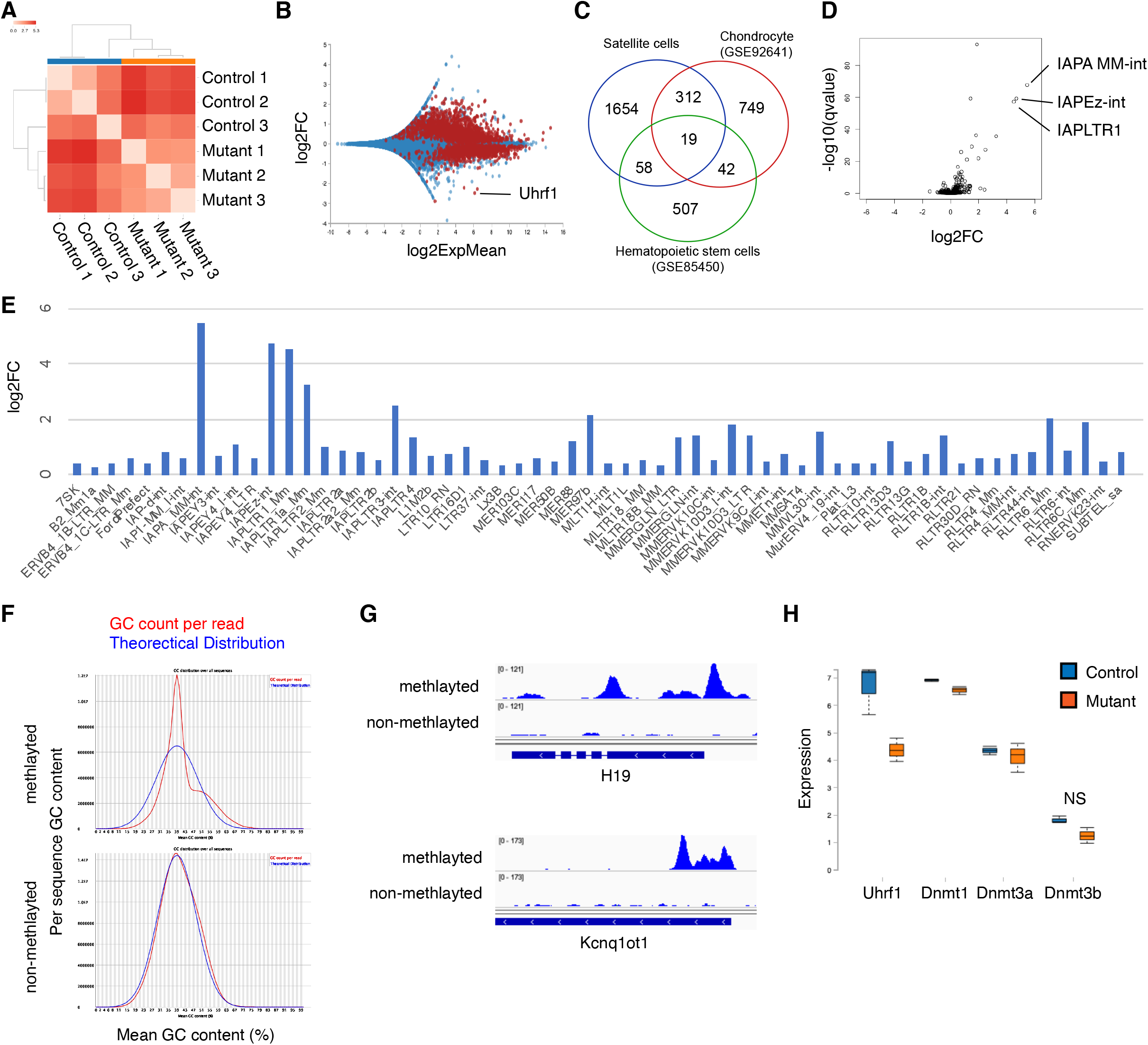
Loss of Uhrf1 in satellite cells induces a distinct gene expression profile, including transposable elements (TEs), and alters the DNA methylation pattern. (A) Heatmap of the expression similarities among samples. (B) MA of the differential expression results. Genes with a significant change in expression are highlighted as red dots. (C) Venn Diagram showing the genes commonly up-regulated by deletion of Uhrf1 among satellite cells, chondrocytes (GSE92641), and hematopoietic stem cells (GSE85450). (D) MA of differential expression of TEs. (E) Up-regulated TEs in mutant satellite cells. (F) GC content per read of methylated and non-methylated DNA extracted from control satellite cells. (G) Distinct methylation patterns observed in H19 and Kcnq1ot1 genes in methylated and non-methylated DNA from control satellite cells. (H) Boxplot showing the expression distribution (normalized as TPMs) of DNA methylation-related genes in each group.

**Supplementary Figure 4.**
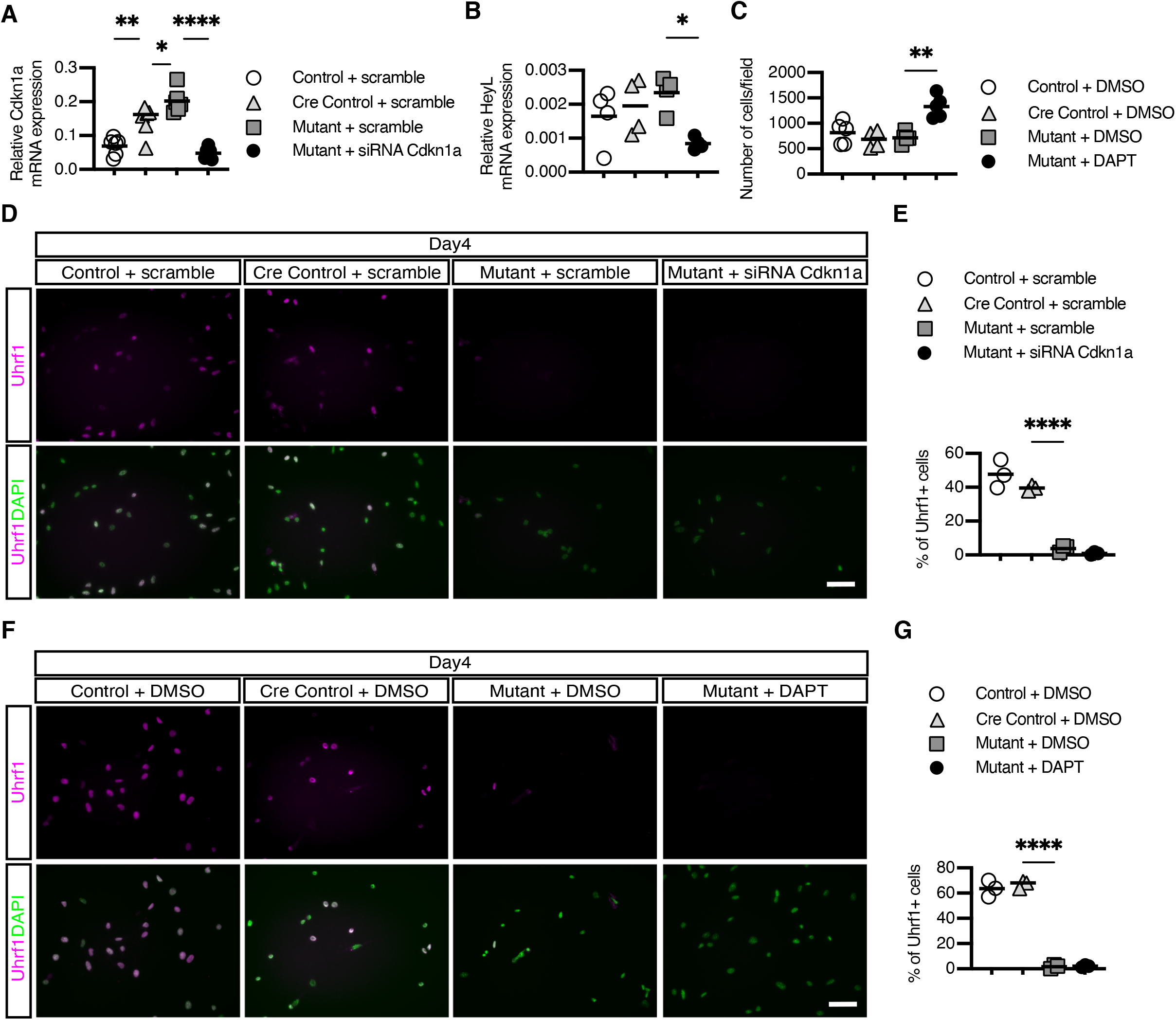
Uhrf1 expression in satellite cells is not altered by knockdown of Cdkn1a or DAPT treatment. (A) RT-qPCR analysis of *Cdkn1a* in satellite cells from control and mutant mice treated with *Cdkn1a* siRNA (n = 6 mice/condition). (B) RT-qPCR analysis of *HeyL* in satellite cells from control and mutant mice treated with DAPT (n = 4/condition). (C) Quantification of the number of cells treated with DAPT (n = 5 for control and mutant; n = 4 for Cre control cells). (D) Immunostaining of Uhrf1 in *Cdkn1a*-knockdown satellite cells. (E) Quantification of Uhrf1+ *Cdkn1a*-knockdown satellite cells (n = 3 mice/condition). (F) Immunostaining of Uhrf1 in satellite cells treated with DAPT. (G) Quantification of Uhrf1+ DAPT-treated satellite cells (n = 3 mice/condition). One-way ANOVA with Šídák’s multiple comparison test was performed, \**p* < 0.05, \*\**p* < 0.01, \*\*\*\**p* < 0.0001. Scale bars = 50 µm.

